# Zeta potential measurements of *Escherichia coli* to evaluate colistin susceptibility and gain insight in resistance mechanisms

**DOI:** 10.64898/2026.01.22.701095

**Authors:** Farras Daffa Imtiyaz, Julien M. Buyck, Luc Deroche, Emma Rose Tewes, Sandrine Marchand, Frédéric Tewes

## Abstract

Colistin resistance in *Escherichia coli* arises from outer membrane (OM) remodeling that reduces surface charge and thereby lowers drug binding affinity. In this study, we investigated the zeta potential of *E. coli* strains with colistin minimum inhibitory concentrations (MICs) ranging from 0.125 to 16 mg/L, following EDTA treatment to chelate divalent cations and unmask intrinsic surface charge. Zeta potential was measured across pH values from 3 to 7, revealing consistent pH-dependent trends and an average 18.5 mV reduction in surface charge with increasing MIC. At pH 7, zeta potential strongly correlated with colistin resistance (R^2^ = 0.919), and this correlation was further strengthened following exposure to sub-inhibitory colistin (1/8 MIC; R^2^ = 0.9975). Notably, resistant strains carrying *mcr-1* or *mcr-4* exhibited significant shifts toward less negative surface charges after sub-MIC exposure, whereas the susceptible parental strain remained unchanged. However, *mcr-1* mRNA expression did not consistently increase under these conditions, highlighting a disconnect between transcriptional responses and phenotypic charge alterations. These findings suggest that regulatory pathways beyond *mcr-1* transcription, including stress-induced lipid A remodeling, contribute to OM charge modulation. Overall, zeta potential profiling provides a rapid and sensitive readout of resistance-associated membrane alterations and complements molecular assays by capturing functional phenotypes. This approach offers a valuable research tool for dissecting colistin resistance mechanisms and may inform future strategies for monitoring bacterial adaptation to last-resort antibiotics.

## INTRODUCTION

Antibiotic resistance is a growing global threat, with bacterial pathogens developing resistance faster than new antibiotics are being discovered. The World Health Organization (WHO) highlighted this issue by prioritizing bacterial pathogens of public health importance for drug development. Among these, carbapenemase-producing *Enterobacterales* are particularly concerning (WHO, 2024), often exhibiting resistance to nearly all available antibiotics. Colistin remains one of the few antibiotics effective for the treatment of infections caused by multidrug-resistant Gram-negative bacteria, classified as a last-resort option when its minimum inhibitory concentration (MIC) is less than 2 mg/L, according to the European Committee on Antimicrobial Susceptibility Testing (EUCAST) breakpoints ^2,3^.

Colistin exerts its antibacterial activity by targeting the bacterial membranes. It binds electrostatically to the negatively charged lipid A of lipopolysaccharides (LPS), thereby destabilizing and damaging the outer membrane (OM) before disrupting the inner membrane, which ultimately leads to cell leakage and death ^4^. In *Escherichia coli*, colistin resistance primarily arises from modifications to lipid A, including the addition of 4-amino-4-deoxy-L-arabinose (*L-*Ara4N) and/or phosphoethanolamine (PEtN) to the phosphate groups, which reduce the membrane’s net negative charge and thereby decrease colistin binding affinity. These modifications are regulated by the chromosomally encoded PmrA/PmrB and PhoP/PhoQ two-component systems, which induce the expression of the *arnT* and *eptA* genes encoding the lipid A transferases that catalyze the addition of L-Ara4N and PEtN, respectively ^5,6^. For example, clinical *E. coli* isolates with mutations in PmrA or PmrB display PEtN-modified lipid A structures and colistin MICs of 8–16 mg/L ^7^. In addition to these chromosomal mechanisms, plasmid-mediated mobilized colistin resistance (*mcr*) genes can also confer resistance by encoding PEtN lipid A transferases that modify lipid A ^8^. For instance, *E. coli* YD626 carrying *mcr-1* exhibits a modified lipid A structure and a colistin MIC of 2 mg/L, which is 16-fold higher than that of its isogenic wild-type strain ^9^. Although resistance conferred by *mcr* genes is generally moderate (MICs of 2–8 mg/L), these genes can facilitate the emergence of higher-level, chromosomally mediated resistance, with MICs reaching up to 64 mg/L in *E. coli* and even higher in *Klebsiella pneumoniae* ^10,11^.

Since colistin resistance in *E. coli* is primarily associated with a reduction in the negative charge on the OM, useful information about both the degree of resistance and whether it stems from lipid A modification pathways. Zeta potential measurement, a parameter that reflects the net surface charge of bacteria relative to the surrounding medium, is an effective and relatively rapid method, requiring only a 24-hour bacterial culture prior to analysis. Although zeta potential can be influenced by various factors, such as pH, ionic strength, and the presence of acidic or basic functional groups on the OM, previous studies have shown that it reliably distinguishes between colistin susceptible and colistin resistant strains ^12,13^.

In this study, we measured the zeta potential of *E. coli* strains exhibiting varying levels of colistin susceptibility, with MICs ranging from 0.125 to 16 mg/L. Zeta potential was assessed at different pH levels following EDTA addition immediately prior to measurement, in order to chelate Mg^2^□ and Ca^2^ □ ions and unmask the underlying net negative charge by removing OM–stabilizing divalent cations. To further explore the mechanistic basis of colistin resistance, we analyzed the relationship between zeta potential and *mcr-1* mRNA expression in *mcr-1*-positive strains exposed to 1/8 of their respective colistin MICs, a sub-inhibitory concentration that may reflect levels encountered during inadequate dosing in patients and could potentially trigger adaptive responses from the bacteria.

Our findings show that zeta potential is a sensitive indicator of phenotypic changes in OM charge that align with differences in colistin susceptibility. While *mcr-1* expression did not consistently scale with MIC, zeta potential measurements revealed a clear reduction in OM negative charge among more resistant strains and after exposure to sub-inhibitory colistin concentrations. This discrepancy suggests that additional factors beyond *mcr-1*, such as other resistance genes, regulatory pathways, or structural modifications, may contribute to OM charge modulation. Collectively, these results highlight zeta potential as a promising complementary tool for assessing colistin resistance and dissecting its molecular underpinnings alongside conventional approaches.

## MATERIALS AND METHODS

### Escherichia coli strains

Four *E. coli* strains with various colistin MICs were used: J53 (KACC 16628, MIC 0.25 mg/L), its *mcr-1* transconjugant J53 *mcr-1* (MIC 4–8 mg/L) ^14^, isolate 184 (*mcr-1*, MIC 8–16 mg/L), and isolate 289 (*mcr-4*, MIC 16 mg/L) ^15,16^.

### Preparation of bacteria for zeta potential measurement

Bacteria were revived from glycerol stock by streaking onto Mueller-Hinton Agar (MHA) plates and incubating overnight at 37°C. A single colony was picked and inoculated into 10 mL of Mueller-Hinton Broth (MHB) in a 50 mL tube and cultured overnight at 37°C with agitation. The resulting culture was centrifuged at 4,000 × g for 10 min at 25°C to pellet the cells. To remove divalent cations from the OM, the bacteria were washed three times with 62.5 µM EDTA. The final suspension was adjusted to an OD600 of 0.3 in 62.5 µM EDTA, with the pH set to 3, 5, or 7 to probe charge variations under different conditions.

To assess the effect of colistin exposure on bacterial zeta potential, bacteria were prepared as above with the addition of colistin at 1/8 of the strain-specific MIC during overnight incubation. After exposure, cells were washed as described and resuspended in 62.5 µM EDTA at pH 7 for zeta potential measurement.

### Zeta potential measurement

The zeta potential of bacterial suspensions was measured using a Zetasizer Nano ZS (Malvern Instruments, France). Bacterial samples were adjusted to an OD600 of 0.3 in 62.5 µM EDTA solutions at pH levels of 3, 5, and 7. Measurements were performed at 25°C in folded capillary cells (DTS1070, Malvern). Each sample (1 mL) was equilibrated for 2 min before measurement. Measurements were performed using “protein” as the material parameter and water as the dispersant. For each condition, three technical replicates of 100 cycles were acquired. Zeta potential values were derived from electrophoretic mobility using the Smoluchowski equation implemented in the instrument software.

### The *mcr-1* expression analysis

Bacteria were grown overnight at 37°C with agitation in the presence or absence of colistin at 1/8 of the MIC, reaching a maximum inoculum of 10^9^ CFU/mL. Bacterial samples were centrifuged at 12,000 × g for 5 minutes, and the resulting pellets were lysed using ZR BashingBead™ Lysis tubes (Zymo Research, France) for 15 minutes under horizontal shaking. Total RNA was extracted from each sample using the Quick-RNA Fungal/Bacterial Kit (Zymo Research, France) following the manufacturer’s instructions. Residual genomic DNA (gDNA) was removed with on-column DNase I treatment (Zymo Research, France) and further treated with the TURBO DNA-Free™ Kit (Thermo Fisher Scientific, France) to ensure complete DNA elimination. RNA concentrations were measured using the NanoDrop One system (Thermo Fisher Scientific, France).

The purified RNA was reverse transcribed into complementary DNA (cDNA) using the FIREScript® RT cDNA Synthesis Kit (Solis BioDyne, Estonia) and an iCycler PCR system (Bio-Rad, France), following the manufacturer’s protocol. Primers targeting the *gapA* reference gene and the *mcr-1* gene were employed as described by Cannatelli et al. (2017) and Bontron et al. (2016).

Quantitative PCR was performed from cDNA with the ONEGreen^®^ FAST qPCR Premix (Ozyme, France) using the CFX96 Touch Real-Time PCR Detection System (Bio-Rad, France). Each sample was analyzed in duplicate, and the relative expression of the *mcr-1* gene was normalized to the expression of the reference gene (*gapA*). The relative differences in mRNA expression levels were determined using the comparative cycle threshold (Ct) method (2^-ΔΔ^Ct). The results were analyzed with the CFX Maestro Software (Bio-Rad).

### Statistical analysis

Zeta potential measurements of *E. coli* strains with varying colistin susceptibilities were analyzed statistically to assess differences among strains. Data were first evaluated for normality using the Shapiro–Wilk test. Depending on the distribution, either one-way ANOVA or the non-parametric Kruskal–Wallis test was applied to compare mean zeta potentials across strains. When significant differences were detected, post-hoc pairwise comparisons were performed using Tukey’s test (for ANOVA) or Dunn’s test (for Kruskal– Wallis) to identify specific strain differences. To examine the relationship between colistin MIC and zeta potential, Spearman’s rank correlation analysis was conducted. All statistical analyses were performed using GraphPad Prism (8.01), and a significance threshold of p < 0.05 was applied.

## RESULTS AND DISCUSSION

### Effect of the pH on *E. coli* Zeta Potential

The zeta potential of *E. coli* strains with varying levels of colistin susceptibility was measured in EDTA solutions across three pH conditions. All strains showed a clear pH-dependent variation, with lower pH values yielding less negative surface charges compared to higher pH levels (**Figure 1**). This effect was particularly pronounced in the highly resistant strain *E. coli* 289 mcr-4, whose average zeta potential shifted from –4.2 mV at pH 7 to +3.9 mV at pH 3. These results align with previous reports describing similar pH effects on zeta potential in Gram-negative bacteria, including *E. coli* ^19^, *Acinetobacter baumannii* ^20^, and *Yersinia enterocolitica* ^21^. Such changes likely reflect alterations in the ionization states of acidic and basic functional groups within the OM, including components such as LPS, phospholipids, and OM proteins. The chosen pH range (3 to 7) encompasses the pKa values of carboxyl moieties, allowing direct observation of their protonation and deprotonation dynamics. These groups contribute to the surface charge and are primarily found in acidic sugars such as glucuronic acid and 3-deoxy-D-manno-octulosonic acid (Kdo), which are located within the core oligosaccharide of LPS and, in the case of glucuronic acid, also within certain O-antigen structures ^22^. Additional contributions arise from exposed carboxylate groups of aspartate and glutamate residues in OM proteins. Within this pH range, the phosphate groups of lipid A are fully deprotonated at their first pKa, around 1.5 to 2, resulting in a monoanionic state ^23^. Near neutral pH, partial deprotonation of the second proton, with a pKa between 6.5 and 7.5, may occur and further influence the net charge. In contrast, amino groups with higher pKa values above 8 remain protonated throughout this range. This pH window is suitable for detecting lipid A phosphate groups, which are the initial binding sites for colistin and targets of chemical modifications such as L Ara4N or PEtN. At the same time, it preserves the integrity of the OM. More extreme pH conditions, below 3 or above 9, are known to destabilize the membrane and compromise bacterial viability ^24,25^. Interestingly, the magnitude of the zeta potential shift in relation to colistin MIC was consistent across the tested pH range, with an average difference of approximately 18.5 mV as MIC increased from 0.25 to 16 mg/L. However, statistically significant differences between all strains were observed only at pH 7; at pH 5 and 3, variations between *E. coli* J53, J53 mcr-1, and 184 mcr-1 were not significant (Kruskal Wallis followed by Dunn’s tests). Based on these findings, subsequent measurements were performed at pH 7.

**Figure 1.**
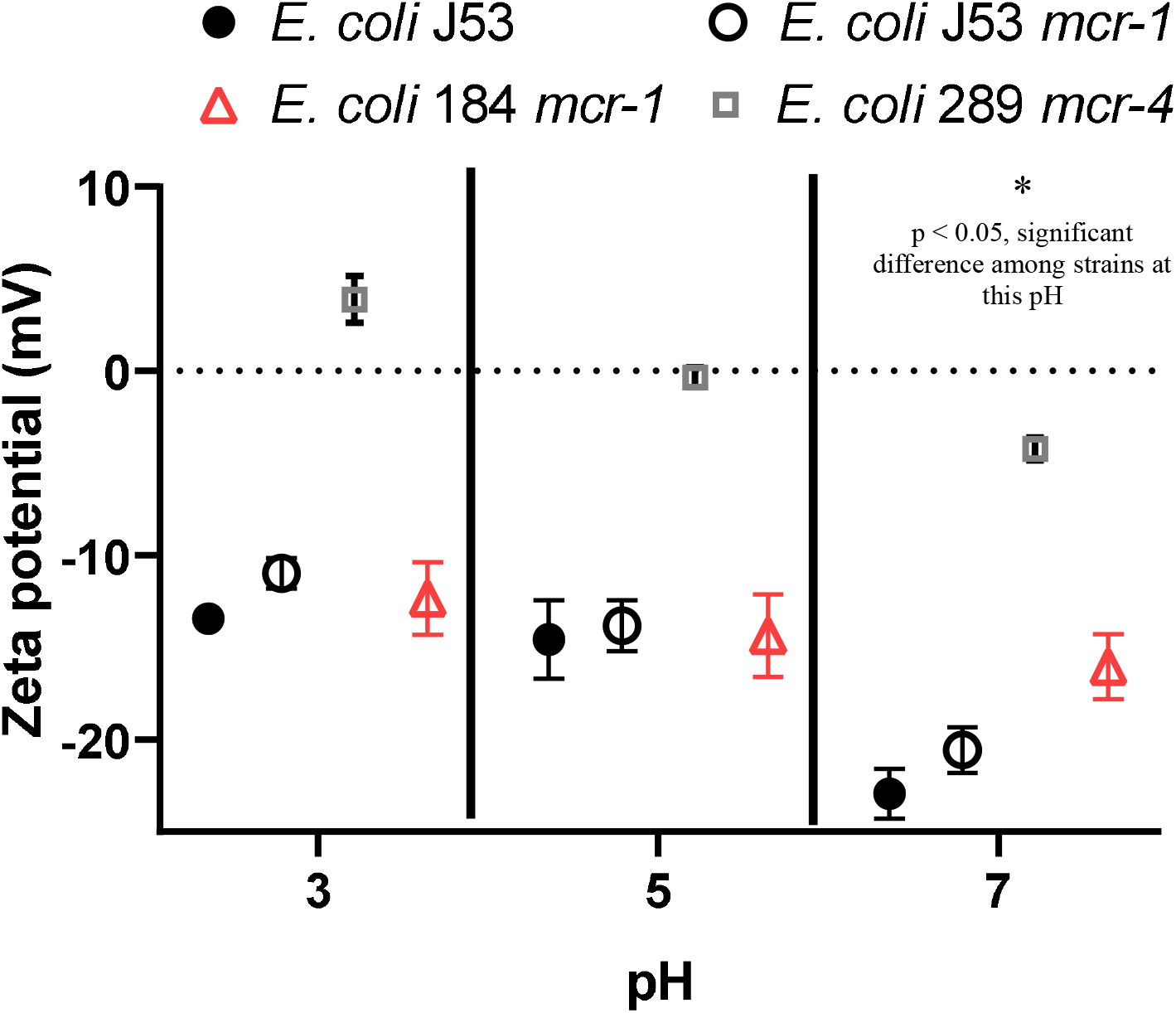
Zeta potential of four *Escherichia coli* strains with varying levels of colistin susceptibility measured across a range of pH values. Bacterial suspensions were prepared in EDTA solutions adjusted to the indicated pH and normalized to an OD □ □ □ of 0.3 before measurement. Values represent mean ± SD from three independent biological replicates performed on separate days. Statistical analysis (one-way ANOVA or Kruskal–Wallis test, as appropriate) revealed a significant difference in zeta potential among strains only at pH 7 (*p* < 0.05).

### Correlation between Zeta potential and colistin resistance in *E. coli* at pH7

At pH 7, zeta potential values were compared with colistin MICs for four *E. coli* strains grown either without or with sub-MIC colistin (1/8 MIC) (**Figure 2**). Across strains, surface charge became progressively less negative as resistance increased, with strong linear correlations observed (R^2^ = 0.919 in untreated strains; R^2^ = 0.9975 after sub-MIC exposure, based on four strains). In untreated cultures, the colistin-susceptible strain J53 displayed the most negative surface charge (-22.9 ± 1.4 mV), whereas the highly resistant *E. coli* 289 mcr-4 exhibited a markedly reduced charge (-4.2 ± 0.6 mV), with intermediate values observed for *mcr-1* strains. These findings indicate that differences between susceptible and moderately resistant strains are relatively modest, but a pronounced inverse relationship emerges at higher resistance levels. Mechanistically, the reduction in negative surface charge among resistant strains seems consistent with *mcr*-mediated OM remodeling, in which PEtN transferases add positively charged ethanolamine groups to lipid A phosphate moieties. This modification reduces the net anionic charge of the membrane and lowers colistin binding affinity ^9,26–28^.

**Figure 2.**
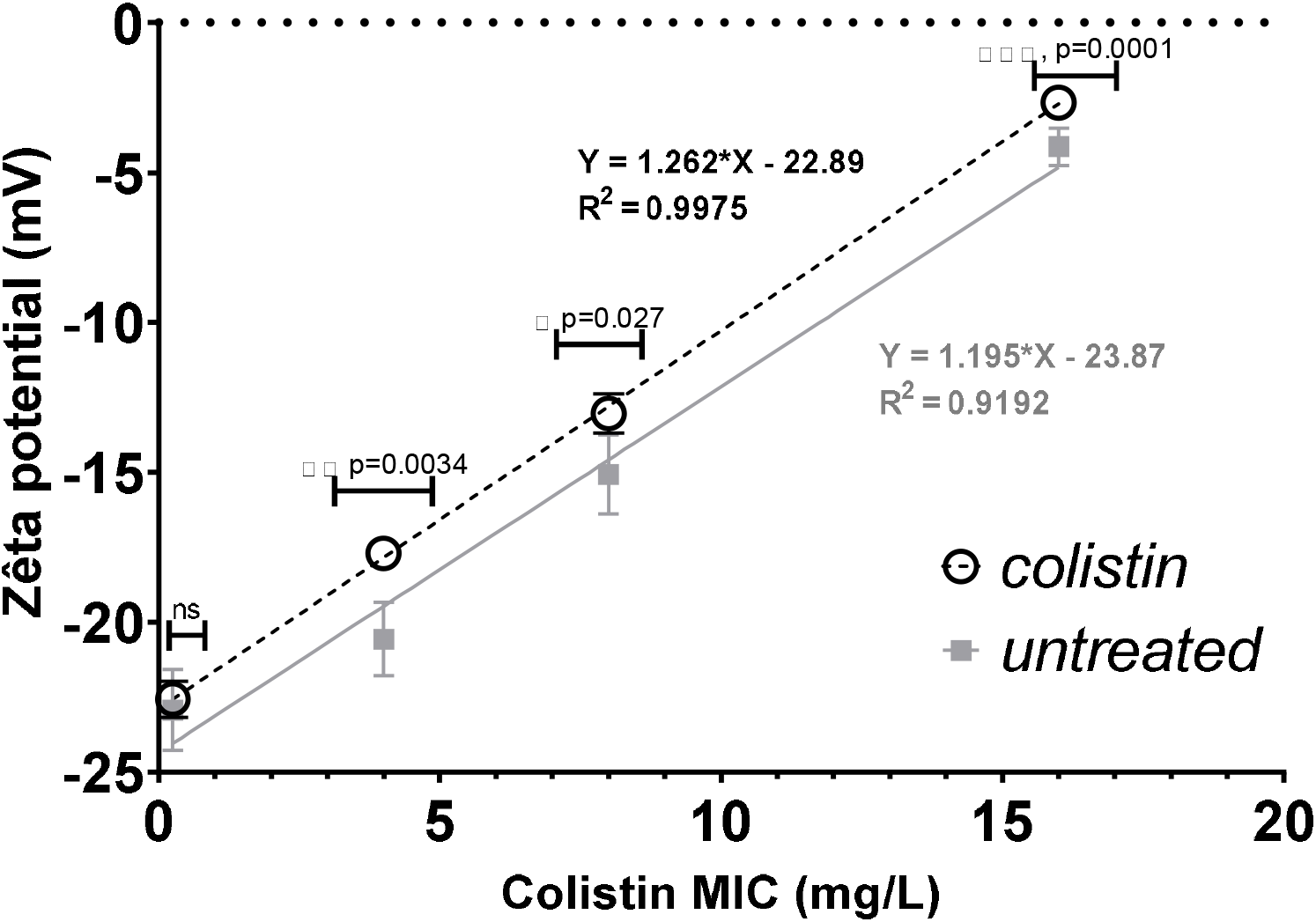
Mean zeta potential of different *E. coli* strains at pH 7 plotted against colistin MIC. Each point represents the mean of three independent measurements performed on three separate days. Solid symbols indicate cultures grown overnight in standard MHB, while open symbols represent cultures grown in MHB supplemented with colistin at 1/8 of the strain-specific MIC. Dotted lines show linear regression of zeta potential versus colistin MIC. Statistical significance between untreated and colistin-treated conditions was determined using a two-tailed Student’s t-test; p-values are shown next to the stars on the graph.

### Effect of sub-MIC colistin on *E. coli* zeta potential and *mcr-1* Expression

Exposure to sub-MIC colistin caused a significant shift toward less negative zeta potential in colistin-resistant *E. coli* strains carrying *mcr-1* or *mcr-4* plasmids, while the colistin-susceptible non–*mcr-1* J53 strain remained unaffected (**Figure 2**). Specifically, *E. coli* J53 *mcr-1* (MIC = 4 mg/L) showed an average shift of 1.6 mV, *E. coli* 184 *mcr-1* shifted by 2.0 mV, and *E. coli* 289 *mcr-4* (MIC = 16 mg/L) exhibited a 1.5 mV shift. These changes likely reflect increased activity of MCR-encoded PEtN transferases or L-Ara4N transferases, which add positive charges to lipid A. Alternatively, colistin could partially neutralize negative charges by binding to LPS, as previously observed ^20^, although extensive washing prior to measurements should have minimized this effect. Although the absolute magnitude of zeta potential shifts did not strongly correlate with MIC, the relative shifts expressed as a percentage of the initial zeta potential were 8.4% for *E. coli* J53 *mcr-1*, 13.6% for *E. coli* 184 *mcr-1*, and 35.7% for *E. coli* 289 *mcr-4*. In contrast, the colistin susceptible *E. coli* J53 (MIC = 0.25 mg/L) maintained a stable zeta potential after sub MIC colistin exposure, as indicated by overlapping error bars.

To determine whether sub-MIC colistin exposure activates the *mcr-1* mechanism, *mcr-1* expression was quantified using real-time PCR in the two *mcr-1*-expressing *E. coli* strains after overnight incubation with sub-MIC colistin. In *E. coli* 184 *mcr-1*, treatment with sub-MIC colistin resulted in a 27-fold increase in *mcr-1* expression compared to the untreated group (**Figure 3**). In contrast, no significant differences in *mcr-1* expression were observed in *E. coli* J53 *mcr-1* under the same conditions.

**Figure 3.**
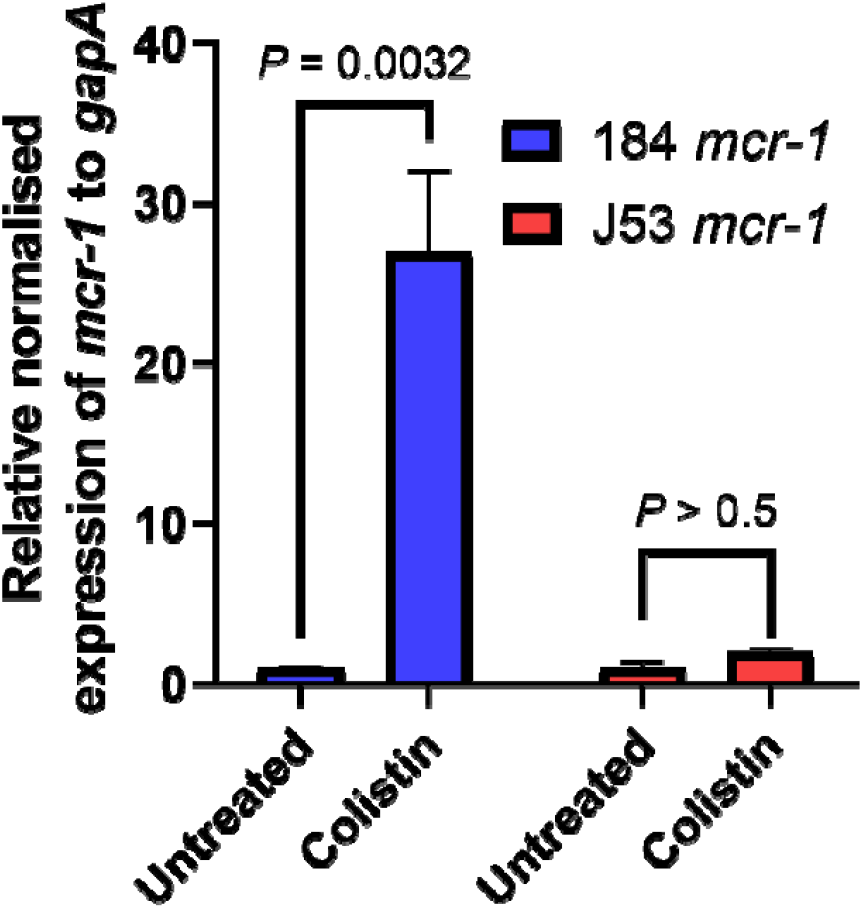
Relative normalized expression of *mcr-1* in *Escherichia coli* strains 184 *mcr-1* and J53 *mcr-1* after overnight culture in Mueller-Hinton broth, with or without supplementation of colistin at 1/8 of each strain’s colistin MIC.

The observation that the zeta potential of *E. coli* J53 *mcr-1* shifted toward less negative values after exposure to sub-MIC colistin, despite no detectable change in *mcr-1* mRNA levels, suggests a disconnect between transcriptional output and phenotypic alterations in membrane charge. This discrepancy may reflect additional regulatory layers. For example, the activity of the MCR-1 phosphoethanolamine transferase can be modulated beyond transcription, through translational efficiency, protein stability, or cofactor availability ^29–31^. Moreover, exposure to sub-MIC colistin is known to activate envelope stress responses, including the alternative sigma factor RpoE and the two-component systems PhoPQ and PmrAB, which drive lipid A and phospholipid remodeling independently of *mcr-1* transcription ^32–34^. Such stress-induced changes in membrane composition provide a plausible explanation for the observed shift in surface charge without corresponding alterations in *mcr-1* expression.

Interestingly, the zeta potential measurement was sensitive enough to detect phenotypic changes following sub-MIC colistin exposure, while RT-qPCR failed to observe a corresponding change in *mcr-1* expression. This emphasizes the complementary nature of these two techniques: RT-qPCR provides insights into transcriptional regulation, whereas zeta potential measurements reveal functional and structural changes at the cell surface. Together, these tools offer a more comprehensive understanding of the bacterial response to sub-MIC colistin, shedding light on both molecular and phenotypic adaptations.

Overall, our results demonstrate that zeta potential is a sensitive indicator of colistin resistance in *E. coli*, capturing both pH-dependent shifts and resistance-associated remodeling of the OM. Importantly, zeta potential measurements complement molecular methods such as RT-qPCR by providing a direct, functional readout of surface charge alterations. While not currently suited for rapid clinical diagnostics, this approach offers a valuable research tool to dissect the multifactorial mechanisms of colistin resistance and could be extended to other OM-targeting agents. Future work integrating zeta potential profiling with statistical modeling and molecular assays may provide a more comprehensive framework for monitoring resistance dynamics.

## DATA AVAILABILITY

The datasets generated and/or analyzed during the current study are available from the corresponding author on reasonable request

## REFERENCES

1. WHO bacterial priority pathogens list, 2024: Bacterial pathogens of public health importance to guide research, development and strategies to prevent and control antimicrobial resistance. https://www.who.int/publications-detail-redirect/9789240093461.

2. Nation, R. L. et al. Framework for optimisation of the clinical use of colistin and polymyxin B: the Prato polymyxin consensus. Lancet Infect. Dis. 15, 225–234 (2015).

3. Pitout Johann D. D., Nordmann Patrice, & Poirel Laurent. Carbapenemase-producing Klebsiella pneumoniae, a key pathogen set for global nosocomial dominance. Antimicrob. Agents Chemother. 59, 5873–5884 (2015).

4. Sabnis, A. et al. Colistin kills bacteria by targeting lipopolysaccharide in the cytoplasmic membrane. eLife 10, e65836 (2021).

5. Rubin, E. J., Herrera, C. M., Crofts, A. A. & Trent, M. S. PmrD is required for modifications to Escherichia coli endotoxin that promote antimicrobial resistance. Antimicrob. Agents Chemother. 59, 2051–2061 (2015).

6. Yan, A., Guan, Z. & Raetz, C. R. H. An undecaprenyl phosphate-aminoarabinose flippase required for polymyxin resistance in Escherichia coli. J. Biol. Chem. 282, 36077–36089 (2007).

7. Sato, T. et al. Contribution of novel amino acid alterations in PmrA or PmrB to colistin resistance in mcr-negative Escherichia coli clinical isolates, including major multidrug-resistant lineages O25b:H4-ST131-H30Rx and non-x. Antimicrob. Agents Chemother. 62, 10.1128/aac.00864-18 (2018).

8. Liu, Y.-Y. et al. Emergence of plasmid-mediated colistin resistance mechanism MCR-1 in animals and human beings in China: a microbiological and molecular biological study. Lancet Infect. Dis. 16, 161–168 (2016).

9. Liu, Y.-Y. et al. Structural Modification of Lipopolysaccharide Conferred by mcr-1 in Gram-Negative ESKAPE Pathogens. Antimicrob. Agents Chemother. 61, 10.1128/aac.00580-17 (2017).

10. Caspar, Y. et al. mcr-1 colistin resistance in ESBL-producing Klebsiella pneumoniae, France. Emerg. Infect. Dis. 23, 874–876 (2017).

11. Zhu, X.-Q. et al. Impact of mcr-1 on the Development of High Level Colistin Resistance in Klebsiella pneumoniae and Escherichia coli. Front. Microbiol. 12, (2021).

12. Esposito Fernanda et al. Detection of colistin-resistant mcr-1-positive Escherichia coli by use of assays based on inhibition by EDTA and zeta potential. J. Clin. Microbiol. 55, 3454–3465 (2017).

13. Hinchliffe, P. et al. Insights into the mechanistic basis of plasmid-mediated colistin resistance from crystal structures of the catalytic domain of MCR-1. Sci. Rep. 7, 39392 (2017).

14. Poirel Laurent, Kieffer Nicolas, & Nordmann Patrice. In vitro study of IS Apl1-mediated mobilization of the colistin resistance gene mcr-1. Antimicrob. Agents Chemother. 61, 10.1128/aac.00127-17 (2017).

15. Al Atya, A.K., Abriouel, H., Kempf, I., Jouy, E., Auclair, E., Vachée, A., Drider, D., 2016. Effects of colistin and bacteriocins combinations on the in vitro growth of Escherichia coli strains from swine origin. Probiotics Antimicrob. Proteins 8, 183–190. 10.1007/s12602-016-9227-9

16. Hazime, N. et al. Enhancing Colistin Activity against Colistin-Resistant Escherichia coli through Combination with Alginate Nanoparticles and Small Molecules. Pharmaceuticals 15, 682 (2022).

17. Cannatelli, A. et al. An allelic variant of the PmrB sensor kinase responsible for colistin resistance in an Escherichia coli strain of clinical origin. Sci. Rep. 7, 5071 (2017).

18. Bontron, S., Poirel, L. & Nordmann, P. Real-time PCR for detection of plasmid-mediated polymyxin resistance (mcr-1) from cultured bacteria and stools. J. Antimicrob. Chemother. 71, 2318–2320 (2016).

19. Hong, Y. & Brown, D. G. Cell surface acid–base properties of Escherichia coli and Bacillus brevis and variation as a function of growth phase, nitrogen source and C:N ratio. Colloids Surf. B Biointerfaces 50, 112–119 (2006).

20. Soon, R. L. et al. Different surface charge of colistin-susceptible and -resistant Acinetobacter baumannii cells measured with zeta potential as a function of growth phase and colistin treatment. J. Antimicrob. Chemother. 66, 126–133 (2011).

21. Schinner, T. et al. Transport of selected bacterial pathogens in agricultural soil and quartz sand. Water Res. 44, 1182–1192 (2010).

22. Klein, G. et al. Molecular and Structural Basis of Inner Core Lipopolysaccharide Alterations in Escherichia coli. J. Biol. Chem. 288, 8111–8127 (2013).

23. Xiao, X., Sankaranarayanan, K. & Khosla, C. Biosynthesis and structure–activity relationships of the lipid a family of glycolipids. Curr. Opin. Chem. Biol. 40, 127–137 (2017).

24. Alakomi, H.-L. et al. Lactic Acid Permeabilizes Gram-Negative Bacteria by Disrupting the Outer Membrane. Appl. Environ. Microbiol. 66, 2001–2005 (2000).

25. Sampathkumar, B., Khachatourians, G. G. & Korber, D. R. High pH during trisodium phosphate treatment causes membrane damage and destruction of Salmonella enterica serovar enteritidis. Appl. Environ. Microbiol. 69, 122–129 (2003).

26. Dortet, L. et al. Optimization of the MALDIxin test for the rapid identification of colistin resistance in Klebsiella pneumoniae using MALDI-TOF MS. J. Antimicrob. Chemother. 75, 110–116 (2020).

27. Furniss, R. C. D. et al. Detection of colistin resistance in Escherichia coli by use of the MALDI Biotyper Sirius mass spectrometry system. J. Clin. Microbiol. 57, 10.1128/jcm.01427-19 (2019).

28. Dortet, L. et al. Rapid detection of colistin resistance in Acinetobacter baumannii using MALDI-TOF-based lipidomics on intact bacteria. Sci. Rep. 8, 16910 (2018).

29. Hu, M. et al. Crystal Structure of Escherichia coli originated MCR-1, a phosphoethanolamine transferase for Colistin Resistance. Sci. Rep. 6, 38793 (2016).

30. Ogunlana, L. et al. Regulatory fine-tuning of mcr-1 increases bacterial fitness and stabilises antibiotic resistance in agricultural settings. ISME J. 17, 2058–2069 (2023).

31. Yang, Q. et al. Balancing mcr-1 expression and bacterial survival is a delicate equilibrium between essential cellular defence mechanisms. Nat. Commun. 8, 2054 (2017).

32. Elizabeth, R. et al. Colistin exposure enhances expression of eptB in colistin-resistant Escherichia coli co-harboring mcr-1. Sci. Rep. 12, 1348 (2022).

33. Huang, J. et al. Regulating polymyxin resistance in Gram-negative bacteria: roles of two-component systems PhoPQ and PmrAB. Future Microbiol. 15, 445–459 (2020).

34. Zeng, X., Hinenoya, A., Guan, Z., Xu, F. & Lin, J. Critical role of the RpoE stress response pathway in polymyxin resistance of Escherichia coli. J. Antimicrob. Chemother. 78, 732–746 (2023).

